# Human Astrocytes Synchronize Neural Organoid Networks

**DOI:** 10.1101/2024.10.17.618921

**Authors:** Megh Dipak Patel, Sailee Sham Lavekar, Ronak Jaisalmeria, Suki Oji, Jazmine Jayasi, Caroline Cvetkovic, Robert Krencik

## Abstract

Biological neural networks exhibit synchronized activity within and across interconnected regions of the central nervous system. Understanding how these coordinated networks are established and maintained may reveal therapeutic targets for neurodegeneration and neuromodulation. Here, we tested the influence of astrocytes upon synchronous network activity using human pluripotent stem cell-derived bioengineered neural organoids. This study revealed that astrocytes significantly increase activity within individual organoids and across long distances among numerous rapidly merged organoids via influencing synapses and bioenergetics. Treatment of amyloid protein inhibited synchronous activity during neurodegeneration, yet this can be rescued by propagating activity from neighboring networks. Altogether, this study identifies critical contributions of human astrocytes to biological neural networks and delivers a rapid, reproducible, and scalable model to investigate long-range functional communication of the nervous system in healthy and disease states.

## Introduction

Biological neural networks are dynamic functional components underlying nervous system activity. In early postnatal development, neural networks have been defined by the presence of synchronous network activity observed in both in vitro and in vivo experimental model systems^1,2^. Synchronized networks can occur in local neighboring neuronal ensembles at the same temporal window, or as spatially propagating waves over long distances. In the adult brain, synchronized oscillatory network activity plays a role in a range of functions such as cognitive control^3^, memory consolidation^4^, and circadian rhythms^5^ that can be dysregulated in various neuropathologies (e.g., schizophrenia, Alzheimer’s disease, epilepsy, etc.). Network activity is primarily synaptically driven and can be experimentally detected as fast synchronized spikes during electrophysiological recordings and as slow oscillatory changes of intracellular calcium, among a population of neurons. However, the extent that which non-neuronal cells and the local environment contributes to this phenomenon is not well established, especially in a human-specific cellular context.

Besides neurons, other cell types, including astrocytes, are thought to dynamically contribute to network connectivity. Astrocytes have multifaceted mechanisms to directly interact and modulate networks including via secretion of various synaptogenic and synaptic adhesion proteins to form and strengthen synapses^6^, bi-directional signaling via neuroactive molecules, homeostatic regulation of ions^7^, production of substrates for neuronal energy metabolism^8,9^, regulation of oxidative stress^10^, and indirect actions via communication with other non-neuronal cells^11^. The culmination of these diverse processes can be dynamically modulated depending on astrocyte activity and reactivity, impacting a wide range of behaviors^12^. Thus, when astrocyte function is perturbed by their reactivity to abnormal states such as disease-associated genetic mutations, environmental insults, and age-related neuroinflammation^13^, then neural networks may be altered by astrocyte-dependent contributions and can potentially be modulated through targeting astrocytes with therapeutics. As we previously reviewed, investigating intercellular contributions of human pluripotent stem cell (hPSC)-derived astrocytes are one promising experimental approach to enable the dissection of specific mechanisms involved in human astrocyte-neural network functional interactions^14,15^.

hPSC-derived neural organoids are emerging tools to model and understand the onset, maturation, and maintenance of neural network activity in baseline and disease-associated contexts. For example, synchronized oscillatory synaptic activity has been observed within single 6-10 month old neural organoids^16^ and between neurons within two organoids physically inter-connected via axon bundles^17^. Altered network activity has also been reported in organoids containing excitatory and inhibitory neurons that have been directly fused together in genetic models of Rett syndrome^18^ and Timothy syndrome^19^. However, because there is an extensive time needed for organoids to exhibit synchronized burst activity in sub-populations of neurons, and because most neural organoid protocols typically result in a mix of multiple cellular subtypes that spontaneously arise over time including progenitor cells, astrocytes, and microglia^20^, the contributions of individual cell subtypes to network activity is obscure and the technology is not well suited for scalable, reproducible high throughput assays of mature networks. Thus, there is a clear need to understand the influence of cells that arise during the onset of neural networks, such as astrocytes, and to optimize the approach for applications ranging from in vitro studies and in vivo engraftments^21-23^.

In this study, we aimed to uncover the contribution of astrocytes to synchronized neural networks using well-defined bioengineered neural organoids composed of post-mitotic excitatory neurons and astrocytes rapidly induced with a transcription factor-based differentiation protocol similar as we previously described^24^. Surprisingly, the presence of astrocytes significantly increases the appearance and frequency of global synchronized activity by as little as 28 days and enables functional connectivity across numerous merged organoids within 24 hours. Further, fusion of healthy organoids to those that cannot display spontaneous synchronized activity due to amyloid treatment can elicit propagation through the degenerating networks, demonstrating a novel assay for screening modifiers of neurodegeneration. Because treatment with the synaptogenic protein, THBS1, and use of specific media components can elicit synchronized bursts in the absence of astrocytes, though at a lesser extent, our study concludes that combinatorial astrocyte support of neuronal health, bioenergetics, and synaptic formation are underlying mechanisms for maintaining persistent synchronized activity.

## Results

### Astrocytes accelerate synchronized burst activity within neural organoids

Our first objective was to determine whether the presence of astrocytes significantly altered the onset and/or characteristics of synchronized brain wave-like global neural network activity throughout human neural organoids. As an experimental readout for synchronized activity, we used neural organoids composed of post-mitotic excitatory neurons that were produced from hPSCs that express a genetic-encoded calcium indicator, GCaMP6, as previously described^24^. At day 1 of neuronal induction via a NGN2 transgene-based approach, organoids were formed in the presence of 20% hPSC-derived inducible astrocytes (aka Asteroids) or in the absence of astrocytes (aka Neurospheres) (**Fig. 1a**), using methods as we previously extensively detailed^24-26^. Before imaging, organoids were stabilized on imaging slides with imaging-compatible media, and then analyzed 16-20 hours later. Surprisingly, by as little as 27-28 days after formation of the organoids, we observed that Asteroids exhibited obvious synchronized oscillations of activity globally throughout the population, indicated by an increased intensity of the calcium indicator throughout the majority of cells (**Fig. 1b, Supplementary movie 1**), yet this was not observed in Neurospheres. We quantified this phenomenon by comparing the mean synchronicity index (MSI)^27^, defined by synchronized change in signal among multiple unbiasedly-selected regions of interest (ROIs) spread out across the entirety of the organoid. The presence of astrocytes resulted in an increased MSI (0.8190±0.04190 for Asteroids and 0.1720±0.04042 for Neurospheres) as well as an increased frequency of synchronized bursts (4.480±0.3600 min^-1^ in Asteroids and 0.5350±0.2267 min^-1^ in Neurospheres) (**Fig. 1c, d**). There was no synchronous network activity observed in either group at 14 or 21 days. This trend was reproduced over 10 independent replicate batches. While these organoids do not display gyrencephalic-like self-organizing structures that are commonly highlighted in organoids generated from long-term culture of progenitor cells, they do exhibit self-organizing structures consisting of soma clustering on one pole of the organoid, reminiscent of a nucleus of neurons in the nervous system, while neuronal fibers with higher noise-to-signal ratios predominantly cluster in the remaining regions (**Extended Data Fig. 1a**). Synchronous burst activity of Asteroids was similarly observed using astrocytes generated from three biologically independent hPSC lines, and were also present with Asteroids that were matured using a small molecule-mediated differentiation protocol^28^, demonstrating that the phenomenon is broadly applicable and is not dependent specifically on an NGN2-based differentiation approach (**Extended Data Fig. 1b, c**).

**Figure 1.**
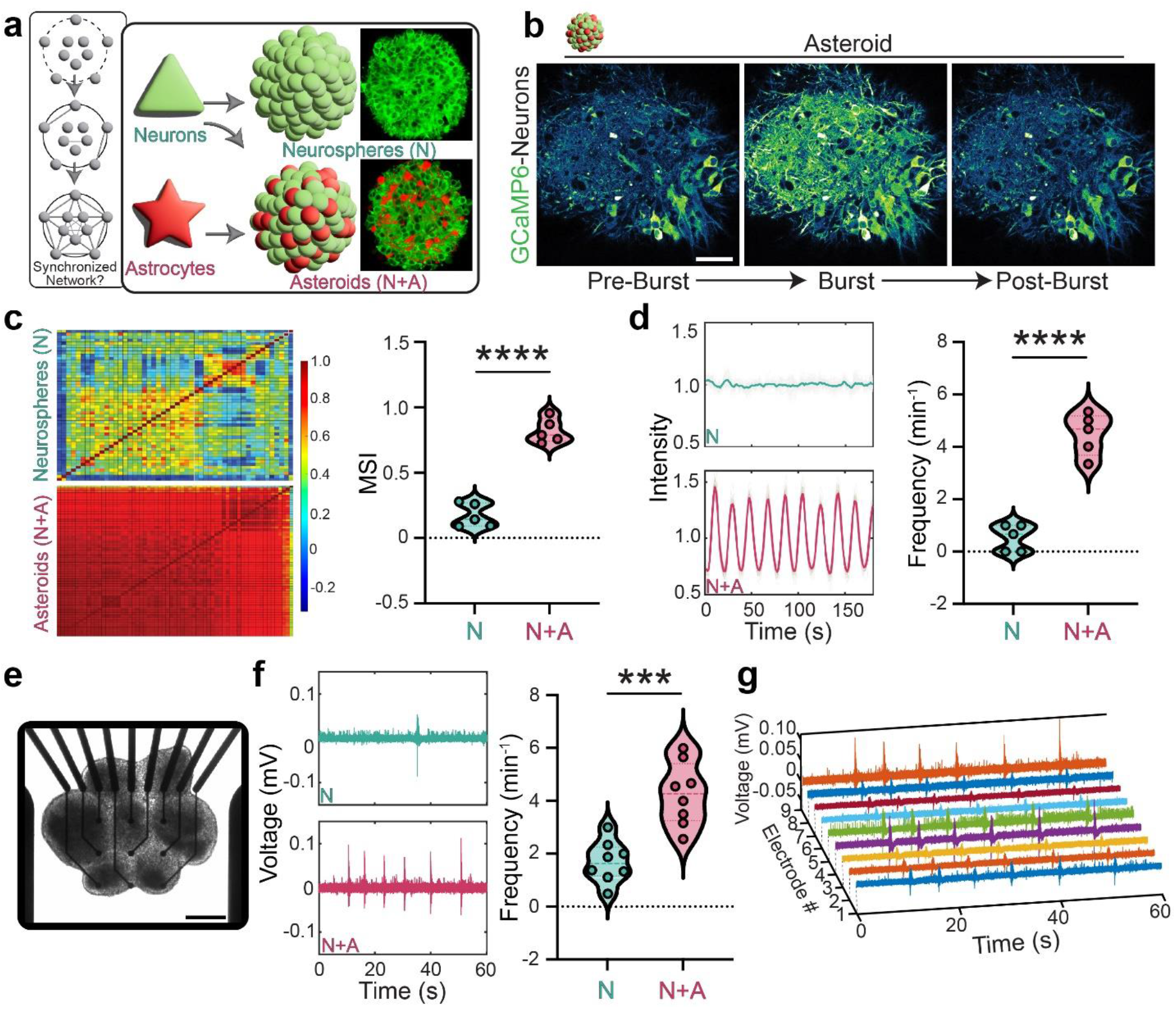
Neural organoids rapidly exhibit synchronous neural network activity in the presence of astrocytes. **a**, Illustrative summary and live image example of neuron-only organoids (Neurospheres) and coculture of neurons and astrocytes (Asteroids). **b**, Asteroids with iGCaMP6-expressing neurons exhibit organoid-wide rise in calcium levels. Scale = 50 µM. **c**, Heatmaps represent the correlation of 64 ROIs during live calcium indicator image acquisition (left) and the resultant mean synchronicity index indicates that Asteroids have increased synchronicity (right) (*n* = 5, *P* < 0.0001, Student’s t-test). **d**, Representative intensity signals over time (left). Asteroids burst with significantly higher frequency compared to Neurospheres (right) *(n* = 5, *P* < 0.0001, Student’s t-test). **e**, Representative live image of Asteroids plated on an array of 3×3 electrodes. Scale: 250 µM. **f**, Asteroids exhibit increased firing of bursts on MEAs when compared to Neurospheres. (*n* = 8, *P* = 0.0002, Student’s t-test) **g**, The bursts observed in Asteroids were synchronous across multiple electrodes spanning several distinct organoids.

Because increases of intracellular calcium are known to correlate with bursts of neuronal activity^29^, we next verified the astrocyte-dependent phenomenon with an alternative approach with higher temporal resolution by using multi-electrode arrays (MEAs), similar as we previously described^24,25^. After 24 hours of incubation of day 35 organoids on MEAs, we recorded extracellular changes of voltage (**Fig 1e**). The presence of astrocytes significantly increased spike burst frequency (4.257±0.4220 min^-1^) (defined by > 5 bursts per 100 msec) compared to organoids in the absence of astrocytes (1.69±0.2746 min^-1^) (**Fig 1f**) at a frequency similar to calcium oscillations. Remarkably, organoids that were physically fused together within 24hr exhibited temporally synchronized burst activity across several spatially distinct organoids in contact with different electrodes, as well as among electrodes in contact with the same organoid (**Fig 1g**), suggesting astrocytes can enable organoids to rapidly form interconnected networks. In summary, this data concludes that astrocytes enable the rapid development of synchronous and robust network activity throughout neural organoids and that Asteroids are an efficient organoid-based experimental model of biological networks.

### Synchronous activity in organoids is dependent on synaptic activity and bioenergetic support

We next aimed to uncover the astrocyte-dependent mechanisms underlying their influence upon synchronous networks. First, we tested whether the astrocytes generated in this study (i.e., those differentiated using a inducible transcription factor-based system) increased synaptic density within organoids similar as we previously observed using astrocytes generated through extensive temporal maturation^25,26^. Indeed, immunostaining of the pre-synaptic marker synaptophysin confirms that the presence of astrocytes leads to an increased synaptic puncta density (5420±816.6 mm^-2^) compared to neuron-only organoids (2099±322.8 mm^-2^) (**Fig 2a**). Additionally, we generated neurons expressing GCaMP8s fused to synaptophysin protein (syp-GCaMP8s) to assess if synchronous calcium oscillations occur locally at synapses. Asteroids with neuronal syp-GCaMP8s were confirmed to similarly display synchronous calcium oscillations among puncta globally (**Extended Data Fig. 2, Supplementary movie 2**). To determine if synchronous network activity is dependent on synaptic activity, we treated Asteroids with antagonists and agonists of neurotransmitters. As a control, we confirmed that addition of 2% volume water in media had no immediate impact. In contrast, acute addition of 2% volume CNQX (10 µM final), an AMPA/Kainate receptor antagonist that blocks excitatory glutamatergic transmission at the synapse, immediately suppressed burst frequency (2.222±0.3742 min^-1^ before CNQX and 0.05550±0.05550 min^-1^ after CNQX) (**Fig. 2b-c, Supplementary movie 3**). Similarly, treatment with the ATP-diphosphohydrolase, apyrase (50 units/mL), blocked burst frequency (pre-apyrase=5.000±0.8074 min^-1^ before apyrase and no detectable activity after apyrase) (**Fig. 2d**). Individual, non-synchronized, activity within single neurons continued to persist in the presence of both CNQX and apyrase, suggesting that network activity is dependent on synaptic transmission without generally disrupting intracellular calcium oscillations. Conversely, to verify that neurotransmitters can induce global activity in the absence of astrocytes, we applied either L-glutamate (500 µM) or ATP (100 µM) to Neurospheres and subsequently observed an immediate rise in intracellular calcium. Treatment with L-glutamate exhibited a rapid rise followed by a slow decrease in calcium (**Fig. 2d**) whereas treatment with ATP causes an immediate rise and fall (**Fig. 2e**).

**Figure 2.**
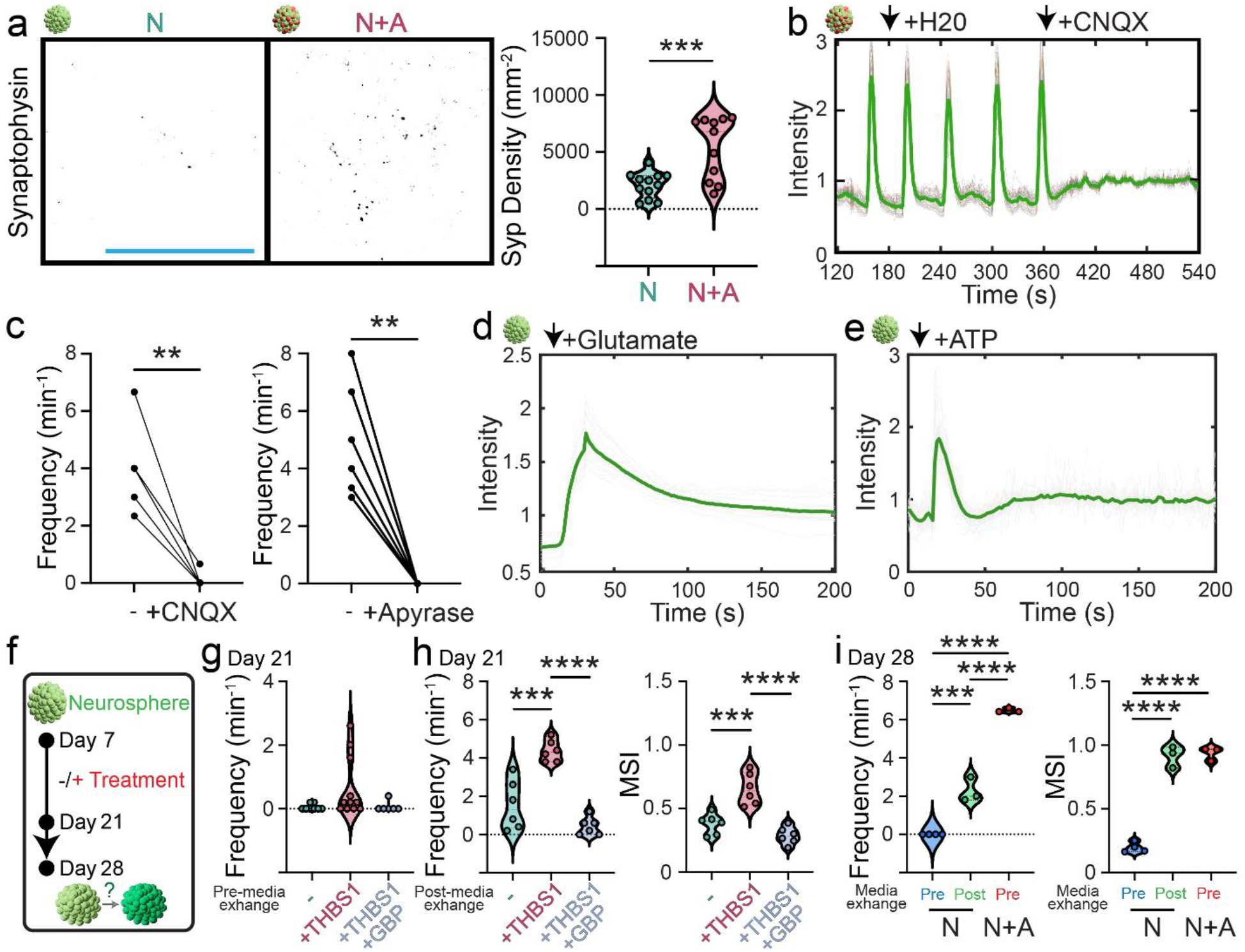
Synchronous activity in Asteroids is mediated by synaptic activity. **a**, Whole-organoid immunohistochemistry staining showing increased density of synaptophysin puncta in Asteroids vs Neurospheres. Scale = 25 µM (*n*_Neurospheres_ = 12, *n*_Asteroids_ = 11 *P* = 0.0008, Student’s t-test) **b**, CNQX treatment results in immediate cessation of burst activity. **c**, CNQX and apyrase treatment upon bursting organoids results significantly lower burst frequencies (*n*_CNQX_ = 6, *P* = 0.0024, Paired t-test; *n*_Apyrase_ = 6, *P* = 0.0016, Paired t-test) **d**, Treatment of L-glutamate onto a non-bursting Neurosphere results in the induction of a single, organoid-wide burst with slow decay. **e**, Treatment of ATP onto a Neurosphere results in a single burst with rapid rise and decay. **f**, Schematic of treatment timeline of Neurospheres. **g**, Neurospheres are unable to burst prior to a partial fresh media exchange. **h**, THBS1-treated organoids exhibit increased MSI and burst frequency compared to no-treatment controls. GBP blocks the effect of THBS1 (*n* = 6, *P*_Frequency_ < 0.0001, *P*_MSI_ < 0.0001, One-way ANOVA). **i**, Neurospheres can exhibit burst activity after a partial fresh media exchange while Asteroids can burst independent of a media exchange. Asteroids burst at a higher frequency than Neurospheres with a media exchange (*n* = 3, *P*_Frequency_ < 0.0001, *P*_MSI_ < 0.0001, One-way ANOVA).

We next sought to test whether astrocyte-mediated synaptogenic factors are sufficient to cause synchronous network activity. We previously determined that hPSC-derived astrocytes secrete significantly larger amounts of synaptogenic proteins compared to neurons, including thrombospondin 1 (THBS1)^24^. THBS1 is a matricellular protein known to activate the α2δ1-receptor on post-synaptic neurons and is blocked by gabapentin (GBP)^30,31^. Thus, Neurospheres were treated with THBS1 protein (3.92 nM)^32^, with or without GBP (32 µM)^33^ starting 6 days after organoid formation (i.e., 7 days after dox-initiated neuronal differentiation) (**Fig. 2f**). At day 21, robust network activity was not observed in all groups (0.05013±0.03282 min^-1^ for untreated Neurospheres, 0.06530±0.2850 min^-1^ for THBS1 and 0.06684±0.06684 min^-1^ for combination THBS1 and GBP treatment) (**Fig 2g**). However, because it is known that media replenishment may induce neural activity^34^, we tested the consequence of a partial exchange with fresh media 30-60 min before measurement. Surprisingly, post media exchange, THBS1-treated Neurospheres exhibited significantly increased synchronous activity at day 21 (Frequency of 4.378±0.2282 min^-1^ in THBS1-treated versus 1.537±0.5272 min^-1^ in untreated Neurospheres and MSI of 0.6548±0.05092 for THBS1-treated versus 0.3782±0.03311 in untreated Neurospheres), and treatment with both THBS1 and GBP led to significantly decreased burst activity (Frequency of 0.4345±0.1897 min^-1^ and MSI of 0.2867±0.02757) (**Fig 2h**). In contrast, Asteroids did not display increased activity at day 21 after media exchange. THBS1-treated Neurospheres also exhibited an increased spike frequency during MEA recordings at 28 days of culture (**Extended Data Fig. 2**). Another known astrocyte-derived regulator of synaptic activity, Glypican 4 (GPC4) (3.92 nM), had no significant influence on burst activity at day 21 (**Extended Data Fig. 2**). Subsequently, we tested the influence of media exchange on organoids at day 28. As previously observed, Asteroids persistently display synchronized activity pre-media exchange (6.484±0.06684 min^-1^ frequency and 0.9363±0.03069 MSI) and Neurospheres do not (0.0000±0.0000 frequency and 0.1973±0.02037 MSI). However, post-media exchange, Neurospheres do exhibit synchronized burst activity (2.273±0.3722 min^-1^ frequency with 0.9163±0.04852 MSI) (**Fig 2i**). Altogether, these studies support the conclusion that astrocytes can accelerate the onset of neural network activity, and enable persistent activity, through a combination of synapse-promoting factors and bioenergetic support.

### Synchronous network burst activity of organoids is a sensitive readout for neurodegeneration and astrocyte neuroprotection assays

During measurement of synchronous activity, we consistently observed a negative correlation between burst frequency and apparent reduced health in a subset of organoid batches that exhibited abnormal morphological characteristics (i.e., loose borders, dark centers, and spontaneously attachment to culture flasks). Because astrocytes are well-known to have neuroprotective attributes^35^, we tested if astrocytes can also underlie robust network activity through enhancing health of organoids during various noxious environmental conditions as a model for neurodegeneration. First, different media conditions were tested because we previously observed that media base and supplemental components can influence organoid viability^24^. Extracellular lactate dehydrogenase (LDH) concentration was used as a general readout for health. LDH secretion after 24 hours in optimal media (i.e., BrainPhys base with complete supplements and growth factors^34^) indicated no difference between Asteroid and Neurosphere health (Asteroids had 100.0±4.147% LDH of Neurosphere control) (**Fig. 3a**). However, Asteroids secreted significantly less LDH after 24 hours in suboptimal media (e.g., DMEM/F12 plus Glutamax without supplements) when compared to Neurospheres (66.31±6.618 % LDH of Neurosphere control), suggesting that astrocytes can contribute to overall improved viability. On the other hand, because inflammatory reactivity of astrocytes has been reported to cause neurotoxicity^13^ and/or dysregulate network activity of hPSC-derived neurons^36^, we tested if this phenomenon occurs in Asteroids by treating with a cocktail of inflammatory cytokines including IL-1α (3 ng/mL), TNFα (30 ng/mL), and C1q (400 ng/mL) from 7 to 28 days of culture. However, inflammation of astrocytes had no effect on network burst activity in either optimal (ITC-treated Asteroids have 146.2±17.72% frequency and 111.6±5.934% MSI of control) or sub-optimal conditions (ITC-treated Asteroids have 130.6±12.17% frequency and 99.54±2.358% MSI of control) (**Fig. 3b**).

**Figure 3.**
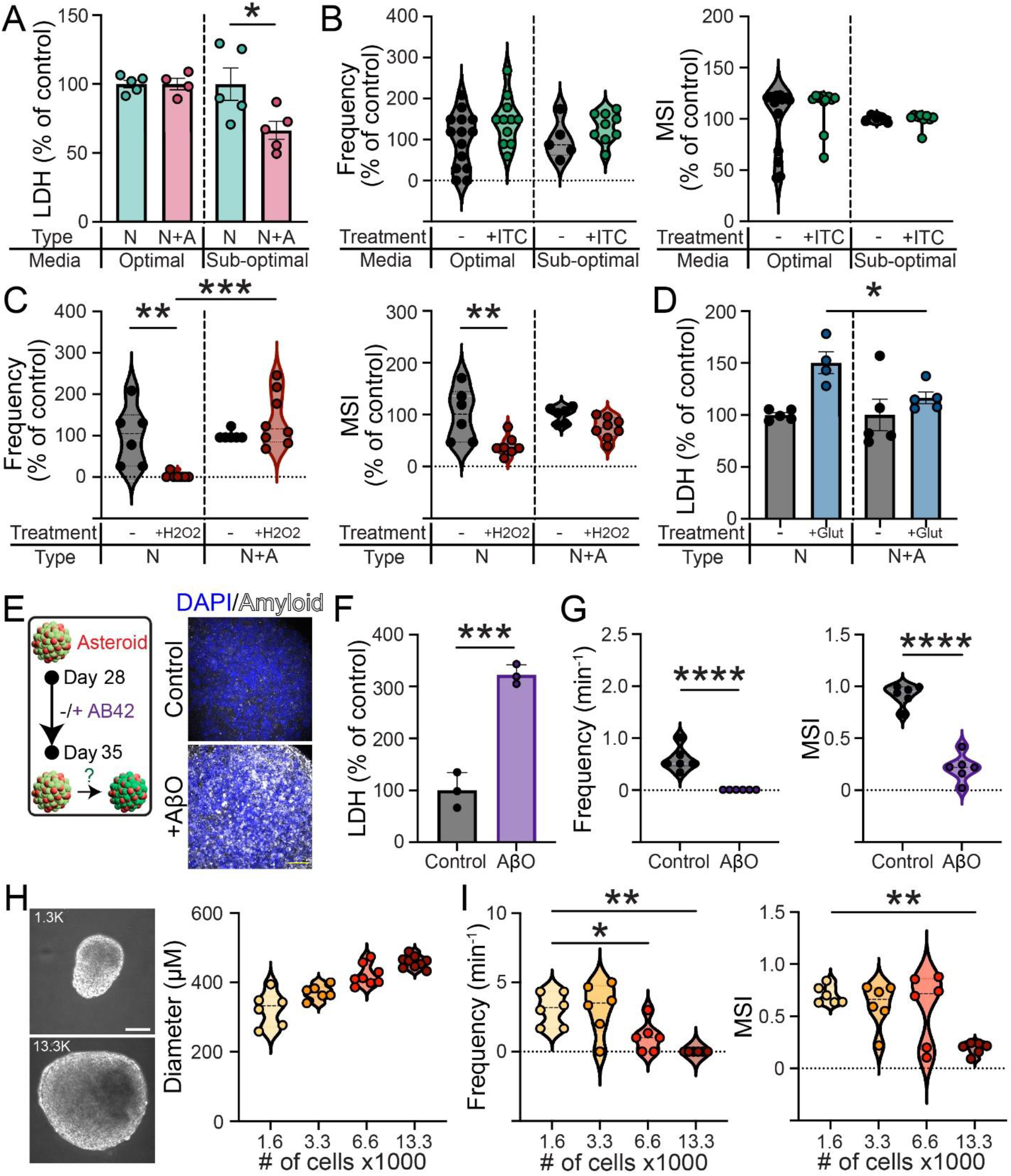
Astrocyte-mediated neuroprotection influences viability and synchronous network activity. **a**, Less LDH is produced in Asteroids compared to Neurospheres when placed in suboptimal media (*n* = 5, *P* = 0.0361, Student’s t-test). **b**, Inflammation of astrocytes does not negatively impact synchronized activity. **c**, Asteroids maintained synchronous network activity when treated with 100 µM H_2_O_2_ while Neurosphere activity is suppressed (*n*_control Neurospheres_ = 6, *n*_H2O2 Neurospheres_ = 7, *n*_control Asteroids_ = 6, *n*_H2O2 Neurospheres_ = 8, *P*_Frequency_ = 0.0002, *P*_MSI_ = 0.0023, One-way ANOVA). **d**, Asteroids treated with 100 µM glutamate secreted significantly less LDH compared to Neurospheres (*n* = 5, *P* = 0.0195, Student’s t-test). **e**, Schematic of amyloid treatment timeline of Asteroids. Amyloid is detected as plaque-like structures throughout the Asteroid. Scale bar = 50 µm. **f**, Treatment of Aβ42 of Asteroids resulted in decreased viability indicated by increased LDH secretion (*n* = 3, *P* = 0.0006, Student’s t-test). **g**, Asteroids treated with Aβ42 exhibit complete suppression of network burst activity (*n* = 6, *P*_Frequency_ < 0.0001, *P*_MSI_ < 0.0001, Student’s t-test). **h**, Asteroids were generated at different sizes by controlling the number of cells in culture. Scale bar = 100 µm. **i**, Asteroids of larger sizes exhibit diminished synchronous network activity (*n* = 6, *P*_Frequency_ < 0.0001, *P*_MSI_ = 0.0002, One-way ANOVA).

To elucidate the role of astrocyte-mediated neuroprotection on network activity, we sought to test individual components that differ between media conditions and are known to affect cell viability, that can also occur via astrocytes, such as protection from oxidative stress and glutamate-induced excitotoxicity. Analysis of previously reported bulk RNA sequencing data, comparing astrocytes to neurons^24^, reveals that astrocytes express significantly more transcripts of antioxidants including superoxide dismutases, glutathione peroxidases, and catalase, suggesting these astrocytes can provide neuroprotection via mitigating oxidative stress (**Extended Data Fig. 3**). To test this, Asteroids were treated with H_2_O_2_ (100 µM) for 24 hours in suboptimal media and this resulted in significant persistence of network burst activity (139.7-±23.25% frequency and 74.11±7.293% MSI of untreated control) in contrast to a dramatic reduction in Neurospheres (2.580±2.580% frequency and 39.72±7.168% MSI of untreated control) (**Fig. 3c**). Importantly, no difference in network activity was observed when both Asteroids and Neurospheres were treated with H_2_O_2_ in optimal media containing antioxidants, suggesting that astrocyte-derived antioxidants may promote improved viability and subsequent network activity (**Extended Data Fig. 3**). Second, RNA sequencing data revealed that astrocytes express significantly more transcripts of proteins that regulate extracellular glutamate levels in comparison to neurons, including glutamate dehydrogenases and glutamate transporters (**Extended Data Fig. 3**). To test if astrocytes protect against glutamate-induced toxicity, 100 µM of glutamate was added for 24 hours and this revealed significantly higher LDH secretion in Neurospheres (150.3±10.52 % LDH of untreated controls) compared to Asteroids (116.5±5.595% LDH of untreated controls) (**Fig 3d**). After treatment, only Asteroids maintained spontaneous activity in neurons (**Extended Fig. 3**), further suggesting that astrocytes can offer protection against the excitotoxic effects of glutamate. As an alternative approach to induce excitotoxicity, we tested astrocyte-mediated protection against neuronal hyperexcitability via optogenetic activation of neurons. Interestingly, blue-light exposure of Neurospheres expressing neuronal ChR2 at 1 Hz pacing for 24 hours revealed a decrease in LDH secretion, suggesting that an increase of neural activity, or blue-light, improves the viability of neural networks. However, optogenetic pacing of Asteroids resulted in no significant difference in viability compared to no-light controls (**Extended Data Fig. 3**).

While oxidative damage and glutamate limit the viability of neural networks and effect network synchronicity, we sought to investigate this phenomenon in response a more specific disease-relevant environmental factor, such as in the context of Alzheimer’s disease (AD)^37,38^. Thus, we treated pre-existing networks in day 28 Asteroids with exogenous amyloid-beta oligomers (Aβ42, 3 µM) for 7 days and analyzed the consequence. Aβ42 resulted in oligomer-laden Asteroids exhibiting a sub-set of structures reminiscent of plaques (**Fig 3e**) and a significant increase in LDH secretion (322.6±11.06% LDH of untreated controls) (**Fig 3f**) indicating a decrease in health. Importantly, treatment of Asteroids with Aβ42 resulted in complete suppression of all spontaneous network burst activity (0.5849±0.09404 min^-1^ frequency with 0.9173±0.04103 MSI in controls compared to 0.0000±0.0000 min^-1^ frequency with 0.2150±0.05282 MSI in Aβ42-treated Asteroids) (**Fig 3g**). Despite no synchronous network activity, Aβ42-treated Asteroids maintained some intraneuronal calcium changes. This is unlike H_2_O_2_ treated Neurospheres, for example, where all intraneuronal spontaneous activity was suppressed. Taken together, this data validates the synchronized neural networks in organoids as a sensitive detector of various disease paradigms.

A major caveat of investigating neural activity in organoids is that larger sizes can be susceptible to central necrosis, leading to reduced viability, as there is limited diffusion of nutrients to the internal tissue. Despite attempts to vascularize neural organoids to mitigate this issue^39,40^, this issue continues to persist and may interfere with the ability of organoids to generate synchronized activity. In Asteroids, we observed that organoids which spontaneously merged into larger aggregates rarely exhibited synchronous network activity. To rigorously confirm this correlation between size and activity, Asteroids of different sizes were created by increasing the number of cells per organoid, ranging from 1.3K to 13.3K (**Fig 3h**). This compares to Asteroids used in prior experiments sized at 3.3K cells per organoid. Calcium imaging revealed that larger Asteroids had a significant decrease in network activity (frequencies of 3.064±0.4913 min^-1^ for 1.3K, 3.119±0.7605 min^-1^ for 3.3K, 1.058±0.4519 min^-1^ for 6.6K, and 0.0000±0.0000 min^-1^ for 13.3K with MSIs of 0.7040±0.03545 for 1.3K, 0.6090±0.08709 for 3.3K, 0.5798±0.1386 for 6.6K, and 0.1933±0.02378 for 13.3K) (**Fig 3i**). This data suggests that organoid size has a significant impact on synchronous networks and this introduce challenges to utilize large organoids that model the multiple interconnected networks across long-distances that occur within the nervous system.

### Astrocytes support long-distance activity propagation through degenerating networks

While physically large spherical Asteroids are unable to rapidly exhibit global synchronous network activity, we observed that spontaneous fusion of several smaller neural networks in a linear manner, reminiscent of an approach sometimes referred to as “assembloids”^41^ to fuse 2-3 organoids together, resulted in spontaneous and synchronous activity among 6 or more organoids within 24 hours (**Fig 4a, Supplementary movie 3**). In contrast, organoids in immediate direct physical contact, did not display this synchronicity, indicating that direct fusion and integration of adjacent organoids is required for this rapid functional interconnectivity. To determine if evoked stimulation can also induce this ensemble-like network activity propagation, we fused one optogenetic Asteroid with several that were not optogenetic and then measured activity during MEA recordings. Indeed, Blue light-induced evoked stimulation of the single organoid propagated synchronous network burst activity across all others (**Fig 4b**).

**Figure 4.**
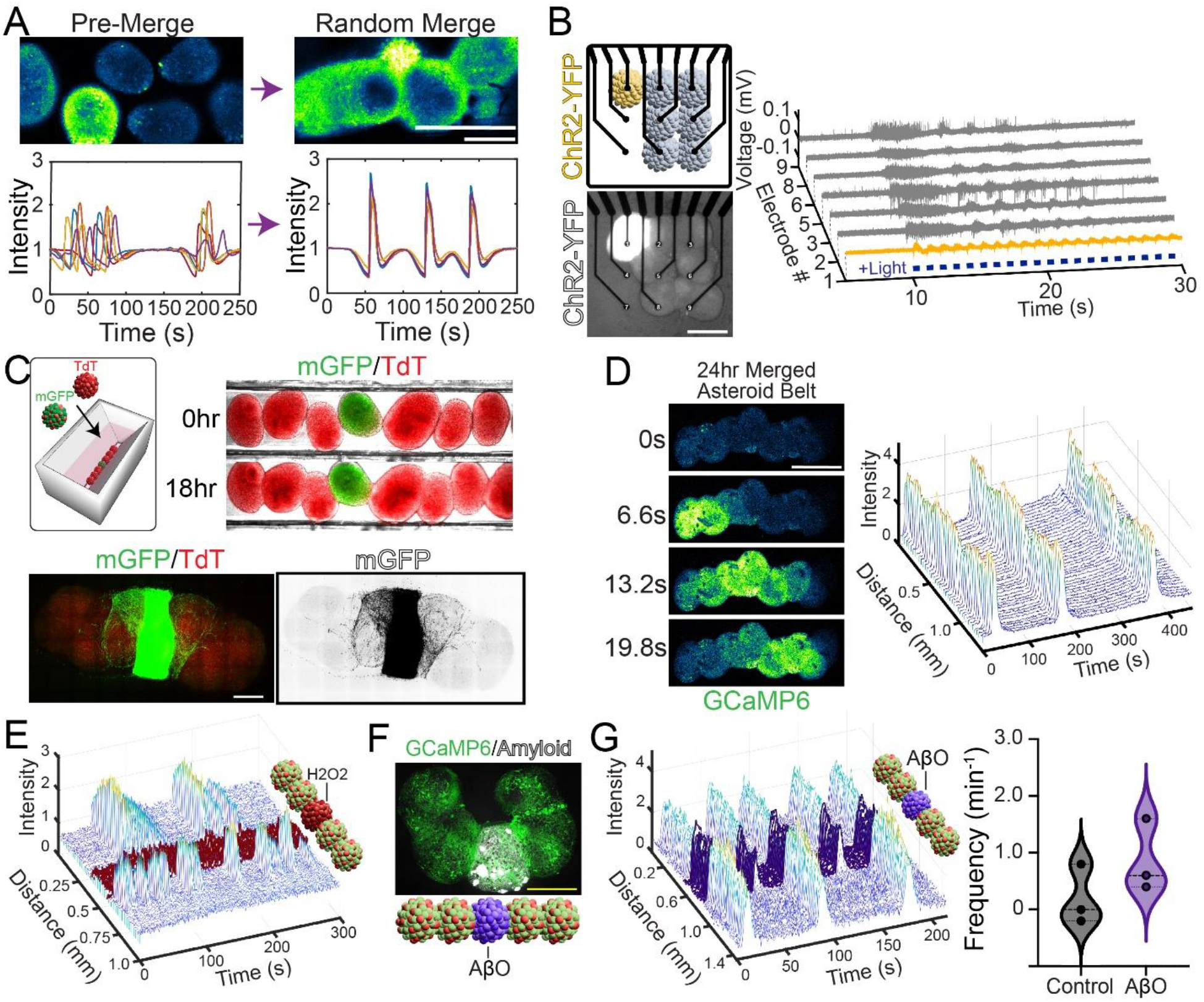
Astrocytes mediate rapid long-range interconnectivity through degenerating neural networks. **a**, Asteroids placed in close-proximity exhibit asynchronous network activity among several organoids, while fusion of the same organoids for 24 hours are synchronous. Scale bar = 250 µm. **b**, Optogetic-evoked activity can propagate when fused to several non-sensitive Asteroids. Scale bar = 250 µm. **c**, Asteroids can fuse together in a linear array within a PDMS mold (top) and rapidly exhibit outgrowth extending across 3 organoids. Scale bar = 250 µm. **d**, Linear fusion of Asteroids reveals synchronous network propagation across an entire Asteroid Belt. **e**, H_2_O_2_-treated Asteroids in the middle disrupt propagation of network activity. **f**, can be inserted between healthy Asteroids. Scale bar = 250 µm. **g**, Propagation of synchronize activity can elicit activity within and through β42-treated Asteroids can be enforced to propagate signals from neighboring nontreated Asteroids (left). The frequency of treated Asteroid Belts was not different than the frequency of untreated (right).

To systematically investigate this phenomenon for reproducible generation of these interconnected networks, we 3D printed a mold to create a PDMS cast to form a physical reservoir for fusing organoids (**Fig 4c**), yielding a linear array term an “Asteroid Belt”. We first sought to analyze the extent of physical connection between organoids within the Asteroid Belt. By 18 hours of fusion, neurite outgrowth between organoids was assessed by placing one organoid expressing membrane-bound GFP (mGFP) in neurons between those expressing tdTomato in neurons. This revealed substantial neurite growth across the immediately adjacent organoid, with some neurites extending across three organoids (**Fig 4c**). Calcium imaging of these Asteroid Belts revealed a wave of network burst activity propagating across the entire distance (**Fig 4d**). Notably, the origin of activity appeared to be initiated by random organoids over time. In contrast, fused Neurospheres in the same fashion exhibited poor network interconnectivity with a lower frequency and synchronicity of network activity (**Extended Data Fig. 4**). Thus, similar to activity within single organoids, the presence of astrocytes mediate long-distance network propagation among multiple organoids.

We next utilized this Asteroid Belt approach to test whether long-distance propagation of pre-existing networks can be influenced by a local region of degenerating tissue, or vice versa, as means to model the spatial heterogeneity that can occur in neuropathological disorders such as Alzheimer’s disease at the cellular^42^ to systems^43^ level. Thus, heterogeneous Asteroid Belts were generated by incorporating a single organoid, treated 24hr with a high dose of H_2_O_2_ (2 µM) to completely inhibit viable network activity, between multiple healthy organoids on either side. 18-24 hr after fusion, synchronized activity was observed among the healthy Asteroids. However, the activity was asynchronous on either side of the H_2_O_2_-treated organoid (n=3 independent replicates) (**Fig 4e, Supplementary movie 4**). This further suggests that robust physical connectivity between healthy neighboring networks is required for signal propagation. As a more specific model of neurodegeneration, we utilized Aβ42-treated organoids that do not exhibit global synchronized activity but still maintain some intraneuronal activity. A single 7 day-treated Aβ42-treated Asteroid was inserted in the middle of the Asteroid Belt and the presence of amyloid-positive plaque-like structures were confirmed (**Fig 4f, Supplementary movie 5**). Surprisingly, synchronized network activity was able to propagate through the treated organoid and progress to the other side of the Asteroid Belt (**Fig 4g**). There was no difference in frequency of network bursts between the Aβ42-treated (0.6000±0.1528 min^-1^) and the untreated (0.9333±0.1856 min^-1^) Asteroid Belts. Altogether, this data demonstrates the potential of healthy astrocyte-laden networks in recovering long-distance activity through damaged neural networks and delivers an advanced testing platform to investigate heterogenous networks.

## Discussion

This study uncovers the multifaceted roles of human astrocytes in developing and maintaining neural network activity within organoid cultures as a simplified, yet well-defined, model of the human nervous system. This was enabled via applying several advancements to neural organoids including genetic engineering to take advantage of induced differentiation and genetic encoded tools, bioengineering principles to maintain consistent size and health, and an innovation method for physical fusion to assess long-distance outgrowth and functionality. As an alternative to allowing hPSC-derived neural progenitors to slowly and spontaneously mature into neurons and astrocytes over extensive length of time, the coculture approach described here uses post-mitotic cells to gain insights specifically into the astrocyte-derived mechanisms contributing to excitatory neural network development and maintenance. A similar modular strategy can be used to study the effect of other cell types that are known to affect synaptic networks including microglia, oligodendrocytes and interneurons. Use of genetic disease-associated cell lines in this approach can further reveal relevant phenotypes in local activity and long-range network propagation as a readout for therapeutic testing relevant to network-level function.

The findings from these studies provide evidence that astrocytes are responsible for synchronizing network activity through a combination of mechanisms. First, we confirmed that addition of a single pro-synaptic protein, THBS1, is sufficient to enhance synchronized activity in the absence of astrocytes even earlier than in the presence of astrocytes. However, supra-physiologic doses of THBS1 were used and a media exchange is still required, suggesting that THBS1 is not the sole mechanism of astrocyte-dependent network activity. Astrocytes produce multiple synaptogenic proteins and synaptic activity promoting factors such as lipids that can be tested in this system. On the other hand, astrocytes can also produce synapse-impeding proteins such as SPARC as well as perform synapse elimination^44-47^. Thus, enhancing the influence of astrocytes upon synapses via increasing production of pro-synaptogenic factors, or inhibiting anti-synapse regulators, may be a strategy to enhance network activity in organoids. This study also revealed that astrocytes enable sustained network activity over time without need to exchange media components, suggesting that they can also provide beneficial regulation of ionic homeostasis, osmolarity, and/or bioenergetic substrates that regulate glucose metabolism in neurons (for example, as recently identified in Alzheimer’s disease^48^). Asteroids can be utilized to investigate these diverse functions in the context of neurodevelopment, aging, neurodegeneration, and context-specific astrocyte reactivity within distinct region-specific networks, leading to translational applications. Conversely, they can be used to also uncover the consequence of neuronal activity-dependent influence upon astrocyte function and health.

In addition to providing components to enhance network activity, our findings reveal that astrocytes also protect networks from detrimental factors in the microenvironment including oxidative stress, glutamate, and amyloid oligomers. While it is evident that health of neurons in a network is a prerequisite for coordinated network activity, identifying the specific contributions of human astrocytes for neuroprotection, such as through providing anti-oxidants, can identify promising therapeutic targets for enhanced protection. For example, this may include targeted activation of astrocyte antioxidant response through master regulators such as Nrf2^49^, enhancement of astrocyte glutamate uptake^50^, and increasing astrocyte release of amyloid-degrading enzymes^51^. Modulation of astrocyte inflammation is another potential strategy to indirectly manipulate network activity^52^, yet in these studies we did not observe detrimental effects of inflammatory astrocytes upon network activity. This lack of effect may be due to the lack of additional inflammatory cell types such as microglia and peripheral immune cells or requires additional disease-associated stressors. We expect that astrocyte-dependent neuroprotection can also promote effective engraftment of organoids when implanted into the nervous system for assessing integration in experimental disease models and for the purpose of cellular replacement therapy.

Biological neural networks communicate through long distances that can span multiple brain regions. Organoids specified to different regional identities have been previously reported to model this phenomenon by physically connecting them together from small networks to larger, more complex cellular populations^41,53^. Here, we advance upon this approach by utilizing a simple mold to rapidly fuse numerous Asteroids in a linear arrangement in one axis to maintain short diameters in the other axis for gas and nutrient diffusion. This linear arrangement permits measurements of activity propagation among networks that can be mixed and matched with disease-associated models, region-and/or maturation stage-specific organoids, and with those pre-treatment with unique molecules for drug testing applications. Thus, this delivers a next generation assay focused on long-range functional connectivity. We expect this approach can be multiplexed to design more complex spatial arrangements such as layering Asteroid Belts upon each other in various configurations. This type of large-scale construction of networks has implications to provide complex interconnected networks that exhibit higher processing power needed for future biocomputing and brain-computer interface applications^54^. Finally, as observed in our in vitro studies, it may be possible to transplant numerous small Asteroids within a living nervous system and allowing them to subsequent fuse together for recover long-distance communication or enforce propagation through damaged networks after disruptions from injury and disease.

## Materials and Methods

### Cell maintenance and differentiation

Astrocytes were generated from hPSC lines WTC-11 (Coriell Institute #GM25256), 162D^55^, and 342 (also known as Cellosaurus PPMI.I.1154.2 [CVCL_D5XJ]) similar as previously described^24^. Briefly, hPSCs were cultured on 6-well plates coated with Matrigel (Corning, 83 µg/mL) and fed daily with TeSR-E8 basal medium with supplements (STEMCELL Technologies) and 1x antibiotic-antimycotic and 1x gentamycin (Gibco). To generate induced neurons, hPSCs (line WTC-11) genetically engineered with GCaMP6, mGFP, TdTom, or ChR2, with a *tet*-on NGN2 overexpression vector in the AAVS1 locus^24^, were grown to 100% confluency before adding 2 µg/mL of doxycycline in neural media (NM) consisting of DMEM/F12 + Glutamax, 1x antibiotic-antimycotic, 0.5x B-27 and 0.5x N-2 supplements (Gibco) to begin neuronal differentiation. Similarly, to generate astrocytes, hPSCs (WTC-11, 162D, and 342) with a *tet*-on SOX9 and NFIA overexpression vector were grown to 30% confluency and then treated with small molecules SB43152 (STEMCELL Technologies; 2 µM), DMH1 (STEMCELL Technologies; 2 µM), and XAV939 (STEMCELL Technologies; 2 µM) were added in NM for 6 days. Doxycycline was added starting at day 0 and maintained throughout culture to induce differentiation.

Neurons were alternatively differentiated using modified version of a small-molecule (GENtoniK) rapid differentiation protocol^28^. hPSCs were plated at 25% confluency into NM plus 2 µM each of SB43152, DMH1, and XAV939 for 10 days with removal of XAV939 after 3 days. At this point, SB43152 and DMH1 were removed, and cells were cultured in NM only for an additional 10 days.

### Organoid fabrication and culture

On day 1 and day 12 of doxycycline treatment for neuron and astrocyte differentiation respectively, cells were split using Accutase (Gibco) and counted and plated into microwell plates (STEMCELL Technologies) with 1×10^6^ neurons/µWell for Neurospheres and 8×10^5^ neurons/µWell with 2×10^6^ astrocytes/µWell for Asteroids. The microwell plate were spun down at 300g for 30 seconds to allow for the cell suspension to aggregate at the bottom of the microwells. Organoids were then transferred to T25 culture flasks (Nunc) with 4 mL of NM plus doxycycline. The flasks were then placed on a belly dancer orbital shaker to prevent merging of the organoids (IBI Scientific). On day 7 of neuron differentiation, media conditions were switched to BrainPhys plus supplements, BDNF (10 ng/mL), GDNF (10 ng/mL), ascorbic acid (0.2 µg/mL), and doxycycline. The organoids were incubated for an additional 21 days with media changes every 3 days.

Neurons differentiated using the GENtoniK protocol were split as neural progenitor cells on day 20 of differentiation and were similarly plated on microwell plates with and without astrocytes. Organoids were transferred to T25 flasks into BrainPhys plus BDNF, GDNF, and Ascorbic Acid, and 10 µM DAPT. After 7 days, DAPT was removed and media was further supplemented with rapid induction small molecules including GSK297955 (1 µM), EPZ-5676 (1 µM), Bay K 8644 (1 µM), and NMDA (1 µM). Organoids were fed with this media every 3 days for 7 days.

A 3D-printed mold (CuraMaker) was created to generate a PDMS chamber that allows organoids to settle into a linear channel 300-500 µm wide and 3 cm long. Organoids aged > 28 days are placed into the chamber holding image optimized BrainPhys plus BDNF, GDNF, and Ascorbic Acid and arranged into a line of 5-8 organoids. The organoids are then incubated to fuse over 18-24 hours.

### Calcium Imaging

#### Organoid Imaging

Intact single organoids were plated on 8-well µ-Slides (IBIDI) coated in matrigel (249 µg/mL) in image-optimized BrainPhys plus supplements. The organoids were incubated for 18-24 hours to allow for attachment and stability of the organoids onto the slide. Confocal live imaging was performed using a TCS SPE microscope and a DFC365 FX camera on a HC PL APO 40x/1.30 oil objective (Leica). All live images were taken at room temperature with a duration of 3 minutes at 512×512 resolution at 0.74 frames per second. Images were acquired using the Leica Application Suite X software. Calcium imaging videos were image stabilized using Fiji (National Institutes of Health). In-house java scripts were made to generate unbiased ROIs in an 8×8 grid overlying the entire image. In-house MATLAB (MathWorks) software was used to import calcium waveform data from each of the ROIs and remove ROIs with an average intensity of < 1. The MATLAB script, *Peak Caller*^*27*^, was used to detrend the calcium waveforms and calculate MSI. Frequency of burst activity was performed by additional custom MATLAB software.

Fused organoids were imaged in the PDMS chamber placed on a slide onto the stage of the TCS SPE microscope. Confocal live imaging was performed with a DFC365 FX camera on a N PLAN 5x/0.12 dry objective. All live images were taken at room temperature at 512×512 resolution at 1.05 frames per second. Calcium imaging videos were image stabilized using Fiji. Custom java scripts were made to generate a vector of ROIs. In-house MATLAB software as well as *Peak Caller* was used to process calcium waveform data from each of the ROIs similar to processing of single organoid ROIs. These ROIs were used to generate waveform plots using additional in-house MATLAB software. Frequency of burst activity was defined by the number of bursts of > 4 consecutive organoids bursting together over the duration of the video taken. MSI was calculated by inputting waveforms averaging calcium activity across the entire surface of each organoid in the ensemble in *Peak Caller*.

### Multielectrode Array

Organoids were plated onto 6-well multi-electrode arrays (Multi Channel Systems) coated with matrigel (249 µg/mL) and were incubated for 24 hours in BrainPhys media plus supplements. MEAs were placed onto a multielectrode array system (MEA2100; Multi Channel Systems) fitted with a temperature-controlled stage. Spontaneous activity was measured for a duration of 1 minute for 2 trials and was repeated 24 hours later. In-house MATLAB software was generated to analyze spike frequency by calculating a threshold defined by 5x the standard deviation of the noise per electrode recording. Electrodes with calculated spike frequency of 0 Hz were removed from analysis. Burst frequency calculation was performed by measuring the spike frequency in moving 100 msec windows increasing by 1 msec; windows with a frequency of > 50 Hz were considered bursts. A minimum peak distance of 10,000 was used when calculating number of bursts on the frequency waveform to avoid overestimation of burst frequency. The maximum burst frequency between 2 trials over 2 days was used for analysis.

### Optogenetics

Blue-light LEDs (Thorlabs) were fitted to fiber optic cables (Thorlabs) for chronic 24-hour optostimulation at a 1Hz frequency with power density of 0.11 mW/mm^3^. Fiber optic cables were fixed to a 96-well plate using a custom 3D-printed apparatus (CuraMaker) with the line cone covering 5 wells plated with 5-6 organoids each. Optostimulation on organoids plated on MEAs were stimulated at a 1 Hz frequency with a power density of 0.29 mW/mm^3^.

### Amyloid Oligomer Preparation

Aβ42 amyloid protein was resuspended in anhydrous DMSO was subsequently sonicated for 5 minutes at room temperature. Sonicated amyloid protein was further diluted in anhydrous DMSO and vortexed for 15 seconds. Protein was then incubated at 4°C to form oligomers over 24 hours and was added to organoid cultures at a final concentration of 3 µM.

### Viability Assays

The lactate dehydrogenase (LDH) assay was performed by plating 5-6 into a 96-well plate with 100 µL of appropriate media. Organoids were incubated for 24 hours after which 2 uL of conditioned media was collected and diluted into 48 µL of PBS. Samples were then treated with 50 µL of LDH detection reagent (Promega) and were incubated for 1 hour at room temperature. Plates were read using a luminescence detector (Promega Glo-Max).

### Immunohistochemistry

Organoids were fixed with 4% paraformaldehyde for 30 minutes and washed with PBS 3x followed by a wash with PBS + 0.1% Tween-20 for 10 minutes at 4°C. Organoids were then resuspended in organoid washing buffer (PBS + 0.1% Triton X-100 + 0.4% SDS + 2% Goat Serum) and transferred to a 24-well plate. Primary antibodies were added for 24 hours on a shaker at 4°C. Primary antibodies were then removed, and organoids were washed with organoid washing buffer 3x for 1-hour washes on a shaker at 4°C. Secondary antibodies and DAPI were added for 24 hours on a shaker at 4°C and were washed the next day with organoid washing buffer 3x for 1-hour washes on a shaker at 4°C. After washes, all liquid was removed from the organoids as they were resuspended in FUnGI clearing solution^56^. Organoids were incubated in FUnGI for 20 minutes at room temperature and were subsequently loaded onto imaging slides for microscopy.

### Synapse Stain Imaging and Quantification

Organoids cleared and stained for synaptophysin were imaged in 10 µm stacks at 2048×2048 resolution and a z-step size of 0.5 µm. Images were then loaded onto Fiji and flattened into a single plane using the “Z-project” function. The background of the image was eliminated by using the “threshold” function. Images were then cropped into a 50×50 µm square at random. Synapses were then quantified by using the “analyze particles” with size parameters to isolate synaptic puncta. Synaptic density was measured by dividing the total synapse count by the area of the square measured.

### Statistical Analysis

Measurements are presented as mean ± standard error of mean (SEM), unless otherwise noted. The number of sampled units (*n*) are listed in figure legends. Significance was performed using statistical tests including Student’s t-test, paired t-test, and ANOVA using GraphPad Prism (GraphPad Software, Inc.). Errors were calculated using individual organoids or fused organoids. A single sample in MEA recordings refers to the average activity across the 9 electrodes in a single well. Data from experiments using a single biological replicate are averaged across technical replicates; otherwise, experiments across multiple biological replicates compared averages between each biological replicate. Data distribution was assumed to be normal, but this was not formally tested. In RNAseq analysis, genes identified as having significant changes in expression between different cell types were defined by adjusted P_adj_ < 0.05 as calculated using the DESSeq2 R package. In all figures, *, P ≤ 0.05; **, P ≤ 0.01; ***, P ≤ 0.001; ****, P ≤ 0.0001.

## Acknowledgments

Research reported in this publication was supported by National Institute of Neurological Disorders and Stroke grant R01NS12978 and Philanthropic funding from Paula and Rusty Walter and Walter Oil & Gas Corp Endowment at Houston Methodist. The content is solely the responsibility of the authors and does not necessarily represent the official views of the National Institutes of Health.

## Author Contributions

M.D.P., S.S.L., and R.K. designed experiments and analyzed collected data. M.D.P. and R.K. wrote the manuscript. M.D.P., S.S.L, R.J., S.O., and J.J. performed experiments and collected data. C.C., and R.K. provided intellectual content and scientific feedback.

