## Extended Data and Legends for "Human Astrocytes Synchronize Neural Organoid Networks"

**Extended Data (Patel et al.)**

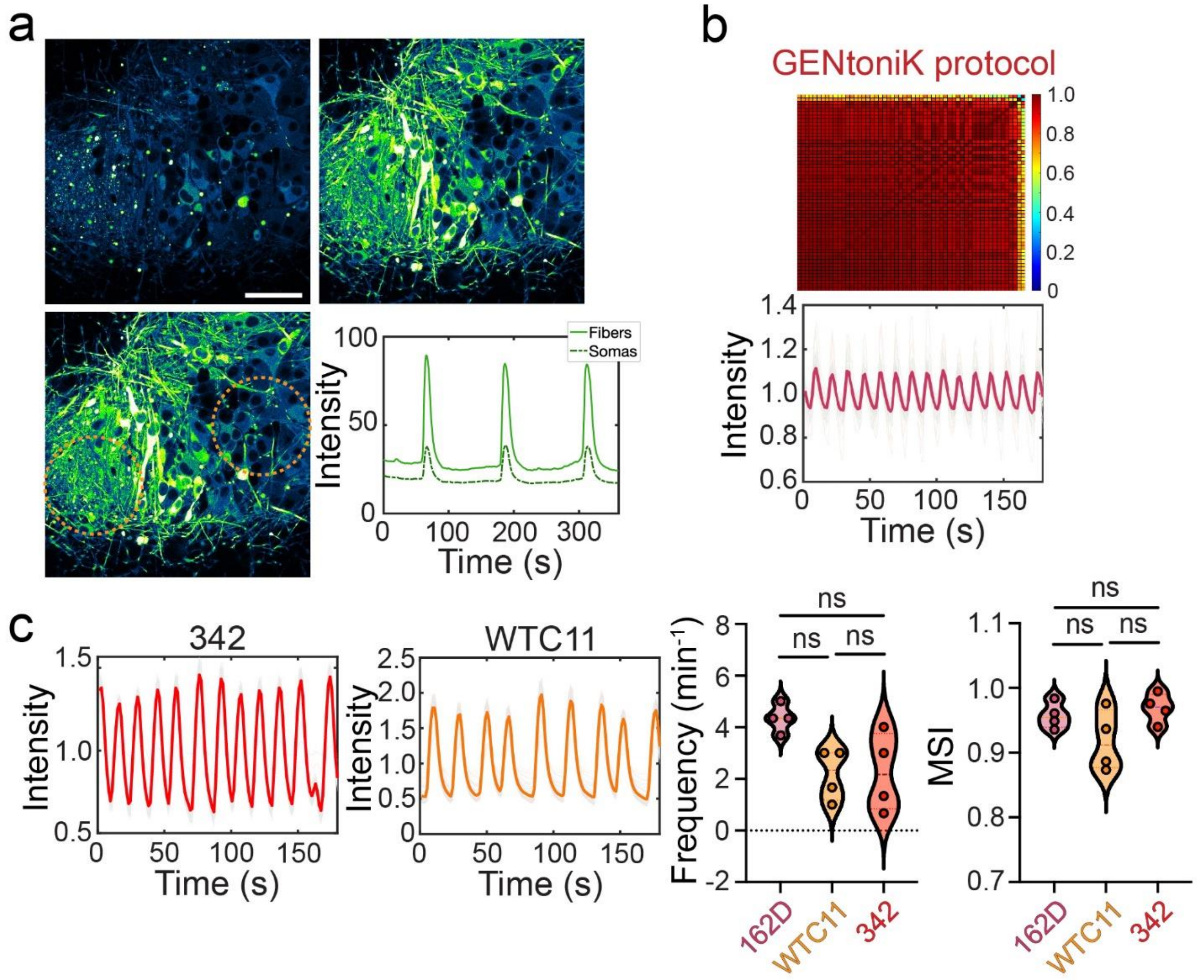


**Extended Data Figure 1. Neural organoids self-organize, and synchronous network activity persists across protocols and biological replicate astrocyte lines. a,** Neural organoids self-organize into fiber-rich and soma-rich regions as they mature with fiber-rich regions exhibiting larger change in calcium bursting network bursts. **b,** Organoids generated with neurons differentiated using the GENtoniK small-molecule cocktail exhibit network burst activity. **c,** Organoids generated using multiple astrocytes differentiated from distinct parental hPSCs reveal similar network burst activity.

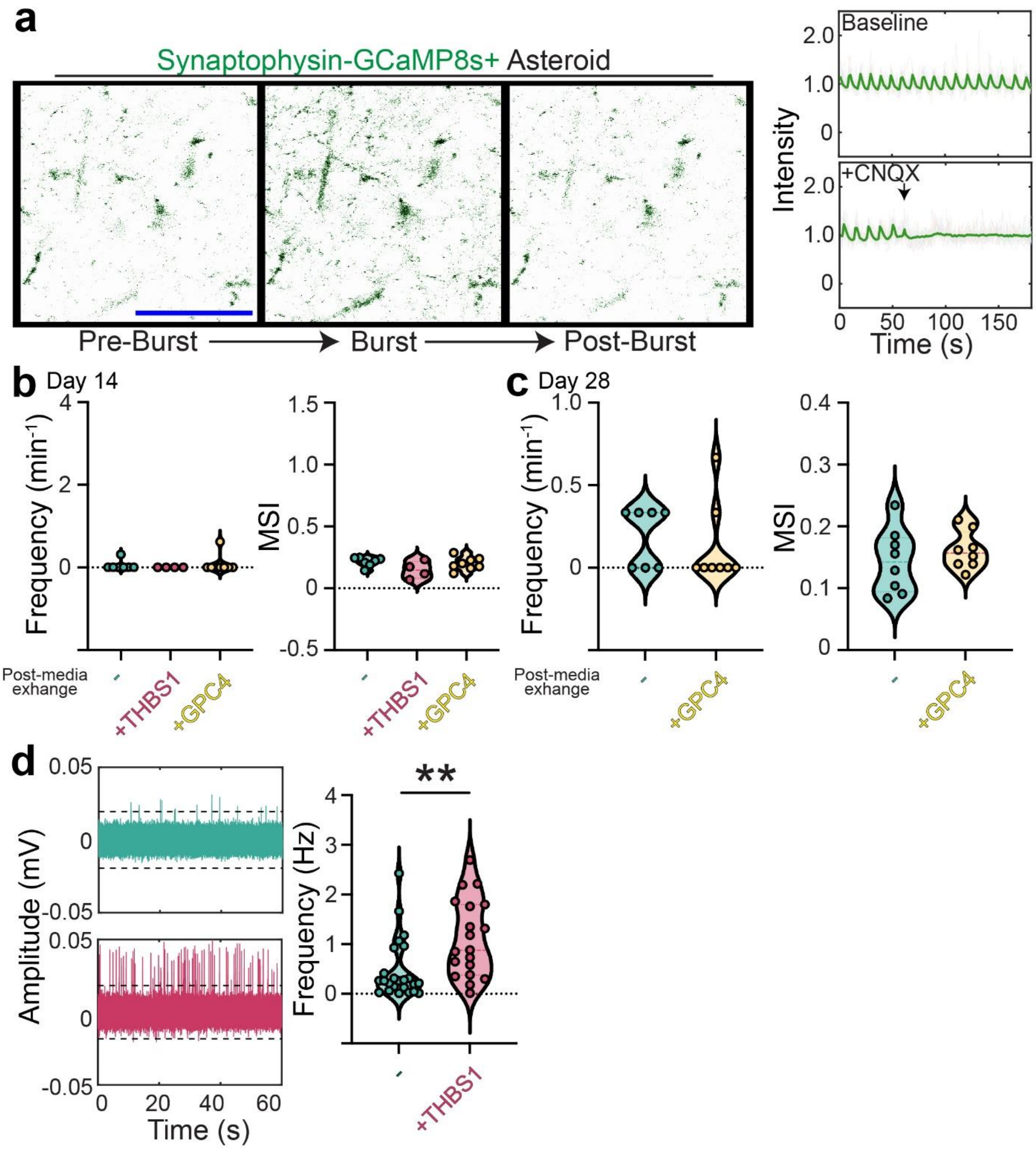


**Extended Data Figure 2. Astrocyte-mediated network burst activity is due to increased synaptic activity. a,** Fluorescent images of a network burst within an Asteroid generated with neurons engineered with synaptophysin-GCaMP8s fusion protein reporter (left). CNQX suppresses of burst activity (right). **b,** Neurospheres treated with THBS1 or GPC4 do not exhibit network burst activity at day 14 despite fresh media exchange. **c,** Neurospheres treated with GPC4 from 7-28 days do not exhibit network activity. **d**, Neurospheres treated with THBS1 results in increased spike frequency of activity on MEAs ($n_{\text{control}}$ = 24, $n_{\text{THBS1}}$ = 19, $P$ = 0.0041, Student's t-test).

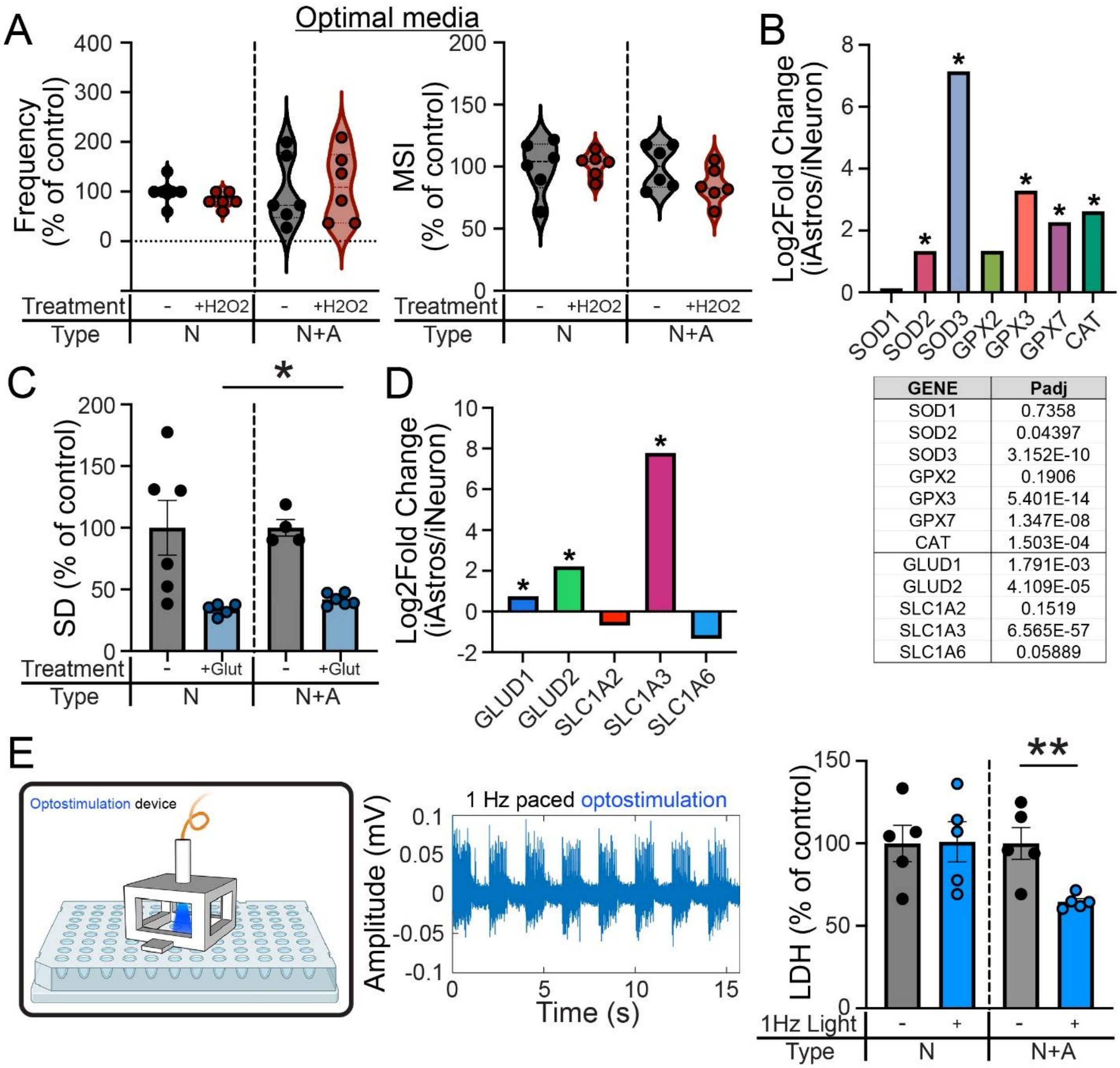


| GENE | Padj |
|---|---|
| SOD1 | 0.7358 |
| SOD2 | 0.04397 |
| SOD3 | 3.152E-10 |
| GPX2 | 0.1906 |
| GPX3 | 5.401E-14 |
| GPX7 | 1.347E-08 |
| CAT | 1.503E-04 |
| GLUD1 | 1.791E-03 |
| GLUD2 | 4.109E-05 |
| SLC1A2 | 0.1519 |
| SLC1A3 | 6.565E-57 |
| SLC1A6 | 0.05889 |

**Extended Data Figure 3. Astrocytes are neuroprotective against oxidative damage and glutamate-induced excitotoxicity, and promotes viability through increased neural activity.** **a,** Asteroids and Neurospheres exhibit no difference in network activity when treated with 100 μM $H_2O_2$ when incubated in optimal media rich in antioxidants. **b,** Astrocytes have significantly higher expression of antioxidant protein-related transcripts. **c,** Asteroids maintain significantly increased intraneuronal activity measured by the standard deviation of calcium changes after treatment of 100 μM of glutamate ($n_{\text{Neurospheres}} = 5$, $n_{\text{Asteroids}} = 6$, $P = 0.0218$, Student's t-test). **d,** Astrocytes have significantly higher expression of glutamate regulatory protein-related transcripts. **e,** Asteroids and Neurospheres generated with neurons engineered to express blue-light sensitive Channelrhodopsin2 can be paced with 1 Hz blue-light. Stimulation of Neurospheres revealed improved viability via decreased LDH secretion with no difference observed in Asteroids ($n = 5$, $P = 0.0072$, Student's t-test).

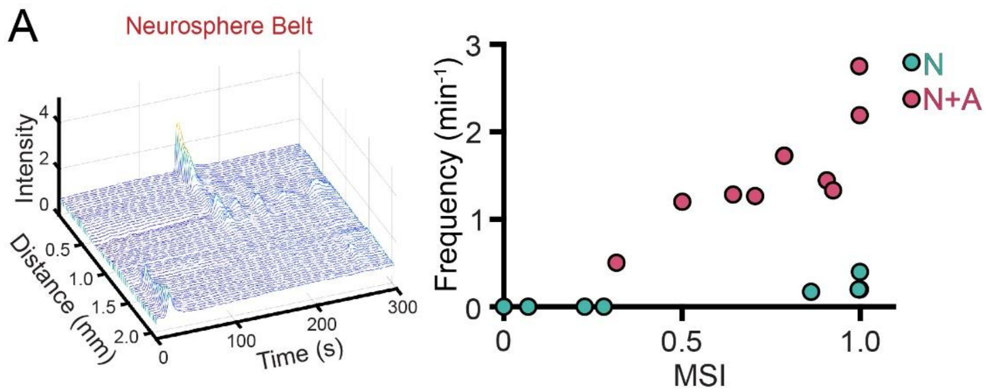


**Extended Data Figure 4. a,** Linearly fused neurospheres rarely exhibit propagation of network activity (left). Scatter plot of frequency of bursts versus MSI reveals that Asteroids burst more synchronously and with higher frequency. ($n_{\text{neurospherebelts}} = 8$, $n_{\text{Asteroidbelts}} = 11$, $P < 0.0001$, Student's t-test) (right).
