## Supplementary Video Legends for "Human Astrocytes Synchronize Neural Organoid Networks"

**Supp. Video 1. Asteroids exhibit synchronous network activity**
Representative example of an Asteroid revealing synchronous calcium activity throughout all neurons in an organoid culture after 28 days of culture. See Fig 1b. Frame rate: 30 fps. Scale bar = 50 μm. Time format = min:sec.

**Supp. Video 2. Asteroids engineered with synaptophysin-GCaMP8s reporter reveal network activity**
Representative example of an Asteroid generated with neurons engineered to express synaptophysin-GCaMP8s fusion protein revealing synchronous calcium activity at the levels of synapses throughout the organoid after 28 days of culture. See Extended Data Fig. 2. Frame rate: 30 fps. Scale bar = 50 μm. Time format = min:sec.

**Supp. Video 3. CNQX selectively blocks synchronous network activity in Asteroids**
Representative example of inhibition of synchronous network activity in Asteroids after addition of AMPA and Kainate receptor antagonist, CNQX. Addition of $H_2O$ does not alter functional network activity. See Fig. 2b. Frame rate: 30 fps. Scale bar = 50 μm. Time format = min:sec.

**Supp. Video 4. Asteroids fused to create an Asteroid Belt exhibit rapid interconnectivity of network activity**
Representative example of eight Asteroids fused to generate an Asteroid Belt revealing propagation of calcium burst activity across the length of Asteroid Belt. See Fig. 4d. Frame rate: 60 fps. Scale bar = 500 μm. Time format = min:sec.

**Supp. Video 5. Dysregulation of network activity using $H_2O_2$ prevents propagation of network activity in Asteroid Belts**
Representative example of an Asteroid Belt featuring a single Asteroid treated with 24-hour treatment of 2 mM $H_2O_2$ revealing asynchronous calcium burst activity among the healthy Asteroids on either side of dysregulated network. See Fig 4e. Frame rate: 30 fps. Scale bar = 500 μm. Time format = min:sec.

**Supp. Video 6. Healthy Asteroids propagate synchronous network activity through AβO-treated networks**
Representative example of an Asteroid Belt featuring a single Asteroid treated with 1-week treatment of 3 μM Aβ42 oligomers revealing calcium activity propagating through the amyloid-treated organoid. See Fig 4f. Frame rate: 30 fps. Scale bar = 500 μm. Time format = min:sec.
